# The origins and consequences of *UPF1* variants in pancreatic adenosquamous carcinoma

**DOI:** 10.1101/2020.08.14.248864

**Authors:** Jacob T. Polaski, Dylan B. Udy, Luisa F. Escobar-Hoyos, Gokce Askan, Steven D. Leach, Andrea Ventura, Ram Kannan, Robert K. Bradley

**Affiliations:** Computational Biology Program, Public Health Sciences Division, Fred Hutchinson Cancer Research Center, Seattle, Washington 98109, USA; Basic Sciences Division, Fred Hutchinson Cancer Research Center, Seattle, Washington 98109, USA; Molecular and Cellular Biology Graduate Program, University of Washington, Seattle, Washington, 98195, USA; David M. Rubenstein Center for Pancreatic Cancer Research, Memorial Sloan Kettering Cancer Center, New York, New York 10065, USA; Human Oncology and Pathogenesis Program, Memorial Sloan Kettering Cancer Center, New York, New York 10065, USA; Department of Pathology, Stony Brook University, New York, New York 11794, USA; Department of Surgery, Memorial Sloan Kettering Cancer Center, New York, New York 10065, USA; Dartmouth Norris Cotton Cancer Center, Lebanon, New Hampshire 03766, USA; Cancer Biology and Genetics Program, Memorial Sloan Kettering Cancer Center, New York, New York 10065

**Keywords:** UPF1, pancreatic adenosquamous carcinoma, cancer genomics

## Abstract

Pancreatic adenosquamous carcinoma (PASC) is a rare and aggressive subtype of pancreatic cancer whose mutational origins are poorly understood. An early study reported somatic mutations in *UPF1*, which encodes a core component of the nonsense-mediated mRNA decay (NMD) pathway, as a common signature of PASC, but subsequent studies did not observe these lesions in other PASC cohorts. The corresponding controversy about whether *UPF1* mutations are important contributors to PASC has been exacerbated by a paucity of functional studies of these lesions. Here, we systematically assessed the potential roles of *UPF1* mutations in PASC. We modeled two reported *UPF1* mutations to find no consistent effects on pancreatic cancer growth, acquisition of adenosquamous features, *UPF1* splicing, UPF1 protein levels, or NMD efficiency. We subsequently discovered that ~40% of *UPF1* mutations reportedly present in PASCs are identical to standing genetic variation in the human population, suggesting that they are likely non-pathogenic inherited variation rather than pathogenic mutations. Our data suggest that *UPF1* is not a common functional driver of PASC and motivate further attempts to identify unique genetic features defining these malignancies.

---

Pancreatic adenosquamous carcinoma (PASC) is a rare and aggressive disease that constitutes 1-4% of pancreatic exocrine tumors^1^. Patient prognosis is extremely poor, with a median survival of eight months^2^. Although PASC is clinically and histologically distinct from the more common disease pancreatic adenocarcinoma, the genetic and molecular origins of PASC’s unique features are unknown.

A recent study reported a potential breakthrough in our understanding of PASC etiology. Liu et al^3^ reported high-frequency mutations affecting *UPF1*, which encodes a core component of the nonsense-mediated mRNA decay (NMD) pathway, in 78% (18 of 23) of PASC patients. These mutations were absent from patient-matched normal pancreatic tissue (0 of 18) and from non-PASC tumors (0 of 29 non-adenosquamous pancreatic carcinomas and 0 of 21 lung squamous cell carcinomas). The authors used a combination of molecular and histological assays to find that the *UPF1* mutations caused *UPF1* missplicing, loss of UPF1 protein, and impaired NMD, resulting in stable expression of aberrant mRNAs containing premature termination codons that would normally be degraded by NMD. The recurrent, PASC-specific, and focal nature of the reported *UPF1* mutations, together with their dramatic effects on NMD activity, suggested that *UPF1* mutations are a key feature of PASC biology.

Three subsequent studies of distinct PASC cohorts, however, did not report somatic mutations in *UPF1*^4-6^. This absence of *UPF1* mutations is significantly different than the high rate reported by Liu et al (0 of 34 total PASC samples from three cohorts^4-6^ vs. 18 of 23 PASC samples from Liu et al; *p* < 10^-8^ by the two-sided binomial proportion test). Although these other studies relied on whole-exome and/or genome sequencing instead of targeted *UPF1* gene sequencing, those technologies yield good coverage of the relevant *UPF1* gene regions because the affected introns are very short. Given this discrepancy, we sought to directly assess the functional contribution of *UPF1* mutations to PASC using a combination of biological and molecular assays.

We first tested the role of the reported *UPF1* mutations during tumorigenesis *in vivo.* Liu et al reported that the majority of *UPF1* mutations caused skipping of *UPF1* exons 10 and 11, disrupting UPF1’s RNA helicase domain that is essential for its NMD activity^7^. We therefore modeled *UPF1* mutation-induced exon skipping by designing paired guide RNAs flanking *Upf1* exons 10 and 11, such that these exons would be deleted upon Cas9 expression (**Fig. 1a**). We chose mouse pancreatic cancer cells (KPC cells: Kras^G12D^; p53^R172H/null^; Pdx1-Cre) as a model system. KPC cells are defined by the *Kras* and *Tp53* mutations that also occur in the vast majority of PASC cases^5,6,8^, making them a genetically appropriate system. We delivered *Upf1*-targeting paired guide RNAs to KPC cells using recombinant adenoviral vectors and confirmed that guide delivery resulted in production of UPF1 mRNA lacking exons 10 and 11 and a corresponding reduction in full-length UPF1 protein levels (**Fig. S1a-c**). We injected subcloned control and *Upf1*-targeted KPC cells into the tails of the pancreata of B6 albino mice (n = 10 mice per treatment) and monitored tumor growth and animal survival. We detected no significant differences in tumor volume or survival in mice implanted with control or *Upf1*-targeted KPC cells (**Fig. 1b-d**). Tumors derived from control as well as *Upf1*-targeted cells displayed similar histopathological features characteristic of moderately to poorly differentiated pancreatic ductal adenocarcinomas (**Fig. S1d**). Moderately differentiated areas were composed of medium to small duct-like structures or tubules with lower mucin production, while poorly differentiated components were characterized by solid sheets or nests of tumor cells with large eosinophilic cytoplasms and large pleomorphic nuclei (**Fig. S1e**). No squamous differentiation was identified by histomorphologic evaluation and no expression of the squamous marker p40 (ΔNp63) was detected (**Fig. S1f-g, Table S1**). We concluded that inducing the reported *Upf1* exon skipping *in vivo* had no detectable effects on pancreatic cancer growth or acquisition of adenosquamous features in the KPC model, although we cannot rule out the possibility that inducing *Upf1* exon skipping in a different model system or cell type could influence tumorigenesis.

**Figure 1.**
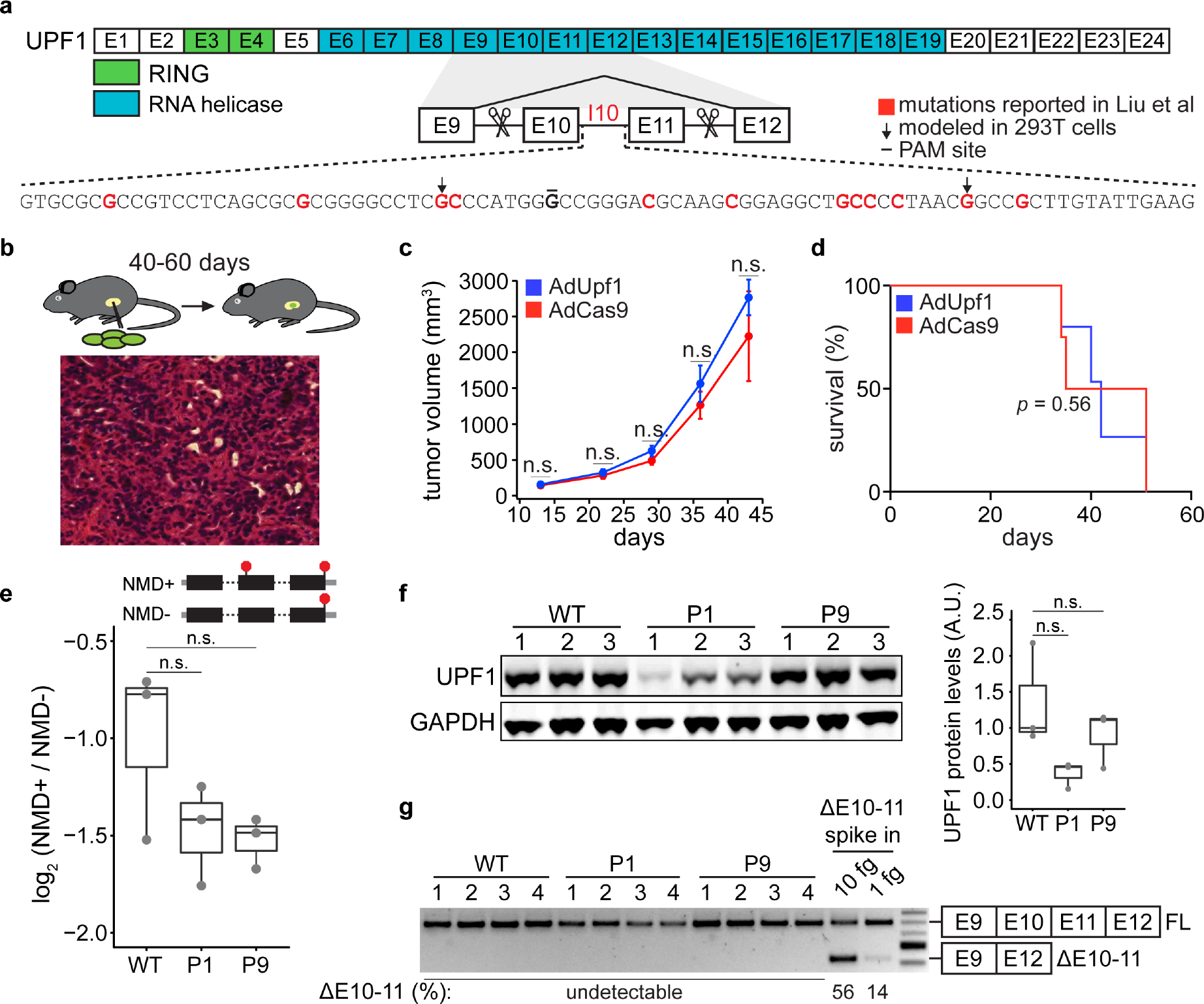
*UPF1* mutations do not confer a growth advantage, alter NMD efficiency, or cause *UPF1* mis-splicing. (**a**) Schematic of *UPF1* gene structure and corresponding encoded protein domains. Intron 10 (I10) contains the bulk of the mutations reported by Liu et al. Scissors indicate the sites targeted by the paired guide RNAs used to excise exons 10 and 11 (E10 and E11). Red nucleotides represent positions subject to point mutations reported in Liu et al. Arrows indicate specific mutations that we modeled in 293T cells. The horizontal black line indicates the nucleotide within the protospacer adjacent motif (PAM) site that we mutated to prevent repeated cutting by Cas9. (**b**) Top, experimental strategy for testing whether mimicking *UPF1* mis-splicing by deleting exons 10 and 11 promoted pancreatic cancer growth. Mice were orthotopically injected with mouse pancreatic cancer cells (KPC cells: Kras^G12D^; p53^R172H/null^; Pdx1-Cre) lacking *Upf1* exons 10 and 11. Bottom, hematoxylin and eosin (H&E) stain of pancreatic tumor tissue harvested from the mice. (**c**) Line graph comparing tumor volume between mice injected with control (AdCas9; Cas9 only) or treatment (AdUpf1; Cas9 with *Upf1*-targeting guide RNAs) KPC cells. Tumor volume measured by ultrasound imaging. Error bars, standard deviation computed over surviving animals (n = 10 at first time point). n.s., not significant (*p* > 0.05). *p*-values at each timepoint were calculated relative to the control group with an unpaired, two-tailed *t*-test. (**d**) Survival curves for the control (AdCas9) or treatment (AdUpf1) cohorts. Error bars, standard deviation computed over biological replicates (n = 10, each group). *p*-value was calculated relative to the control group by a logrank test. (**e**) Box plot of NMD efficiency in 293T cells engineered to contain wild-type (WT) or mutant (P1, P9) *UPF1*. P1 and P9 correspond to the IVS10+31G>A and IVS10-17G>A mutations reported by Liu et al. All cells have the PAM site mutation illustrated in (a). NMD efficiency estimated via the beta-globin reporter assay^11^. Middle line, notches, and whiskers indicate median, first and third quartiles, and range of data. Each point corresponds to a single biological replicate. n.s., not significant (*p* > 0.05). *p*-values were calculated for each variant relative to the control by a two-sided Mann-Whitney U test (*p* = 0.40 for P1, 0.40 for P9). (**f**) Left, immunoblot of full-length UPF1 protein for the cell lines in (e). Each lane represents a single biological replicate with the indicated genotype. GAPDH serves as a loading control. Equal amounts of protein were loaded in each lane (measured by the bicinchoninic acid assay). Right, box plot illustrating UPF1 protein levels relative to GAPDH for each genotype. Middle line, notches, and whiskers indicate median, first and third quartiles, and range of data. Each point corresponds to a single biological replicate. Data was quantified with Fiji (v2.0.0). A.U., arbitrary units. n.s., not significant (*p* > 0.05). *p*-values were calculated for each variant relative to the control by a two-sided Mann-Whitney U test (*p* = 0.10 for P1, 1.0 for P9). (**g**) PCR using primers that amplify both full-length UPF1 mRNA (FL) and mRNA lacking exons 10 and 11 (ΔE10-11). UPF1 mRNA lacking exons 10 and 11 was only detected in the positive control lanes (ΔE10-11 spike in), in which DNA corresponding to UPF1 cDNA lacking exons 10 and 11 was synthesized and added to cDNA libraries created from WT cells prior to PCR. Numbers above each lane indicate biological replicates. Numbers below each lane represent the abundance of the lower band as a percentage of total intensity (see Methods). Data was quantified with Fiji (v2.0.0). See **Fig. S3** for uncropped gels.

We next assessed molecular phenotypes induced with *UPF1* mutations. Liu et al measured the effects of each mutation on *UPF1* splicing using a minigene assay, in which each mutation was introduced into a plasmid containing a small fragment of the *UPF1* gene that was subsequently transfected into 293T cells. Liu et al concluded that all reported *UPF1* mutations caused dramatic *UPF1* mis-splicing that disrupted key protein domains that are essential for UPF1 function in NMD. Minigenes are common tools for studying splicing, but they are frequently spliced less efficiently than endogenous genes, presumably because they are gene fragments that lack potentially important sequence features that promote splicing and incompletely capture the close relationship between chromatin and splicing^9,10^.

We modeled *UPF1* mutations in 293T cells in order to mimic Liu et al’s experimental strategy, but introduced mutations into their endogenous genomic contexts rather than using minigenes. We selected two distinct *UPF1* mutations in intron 10 for these studies. We selected IVS10+31G>A because it was reportedly recurrent across three different patients (making it equally or more common than any other mutation) and induced strong mis-splicing on its own (36% mis-spliced mRNA, versus 0% for wildtype *UPF1*); we selected IVS10-17G>A because it had one of the strongest effects on splicing (90% misspliced mRNA). IVS10+31G>A was present in a homozygous state in two of the three patients carrying it, while IVS10-17G>A was present in a heterozygous state.

We introduced each mutation into its endogenous context by transiently transfecting a plasmid expressing Cas9 and a single guide RNA (sgRNA) targeting *UPF1* intron 10 as well as appropriate donor DNA for homology-directed repair, screened the resulting cells for the desired genotypes, and established clonal lines. The resulting cell lines contained the desired mutations in the correct copy numbers as well as a point mutation disrupting the protospacer adjacent motif (PAM) site (**Fig. 1a, S1h-j**). As neither the PAM site itself nor nearby positions were reported as mutated in Liu et al, we additionally established a cell line in which only the PAM site was mutated as a wild-type control.

We systematically tested the functional consequences of *UPF1* mutations for NMD efficiency, UPF1 protein levels, and *UPF1* splicing. We measured NMD efficiency in our engineered cells using the well-established beta-globin reporter system, which permits controlled measurement of the relative levels of mRNAs that do or do not contain an NMD-inducing premature termination codon, but which are otherwise identical^11^. We did not observe decreased NMD efficiency in *UPF1*-mutant versus wild-type cells; instead, *UPF1*-mutant cells exhibited evidence of modestly more efficient NMD, although these differences were not statistically significant (**Fig. 1e**). Consistent with similar NMD activity independent of *UPF1* mutational status, *UPF1* mutations did not cause loss of full-length UPF1 protein (**Fig. 1f**). Although UPF1 protein levels varied between the individual cell lines, variation in UPF1 protein levels was uncorrelated with NMD efficiency and did not segregate with *UPF1* mutational status. We therefore measured the levels of normally spliced and mis-spliced UPF1 mRNA. We readily detected normally spliced UPF1 mRNA in all samples, but found no evidence of mis-spliced UPF1 mRNA, except in positive control samples in which we spiked in synthesized DNA corresponding to the exon skipping isoform reported in Liu et al (**Fig. 1g**).

Given the differences between Liu et al’s findings of common *UPF1* mutations and their absence from subsequent studies of PASC, we wondered whether some of the *UPF1* mutations reported by Liu et al might correspond to inherited genetic variation rather than somatically acquired mutations. We searched for each mutation reported by Liu et al within databases compiled by the 1000 Genomes Project, NHLBI Exome Sequencing Project, and Exome Aggregation Consortium (ExAC)^12-14^. These databases were constructed from a mix of whole-genome and whole-exome sequencing, both of which are effective for discovering variants within the relevant regions of *UPF1* (because *UPF1* introns 10, 21, and 22 are very short, they are well-covered by exon-capture technologies). We found genetic variants identical to 42.5% (17 of 40) of the reported *UPF1* mutations, one of which is present in the reference human genome. 83.3% (15 of 18) of *UPF1*-mutant patients had one or more reported mutations that corresponded to standing genetic variation (**Fig. 2, S2, Table S2**).

**Figure 2.**
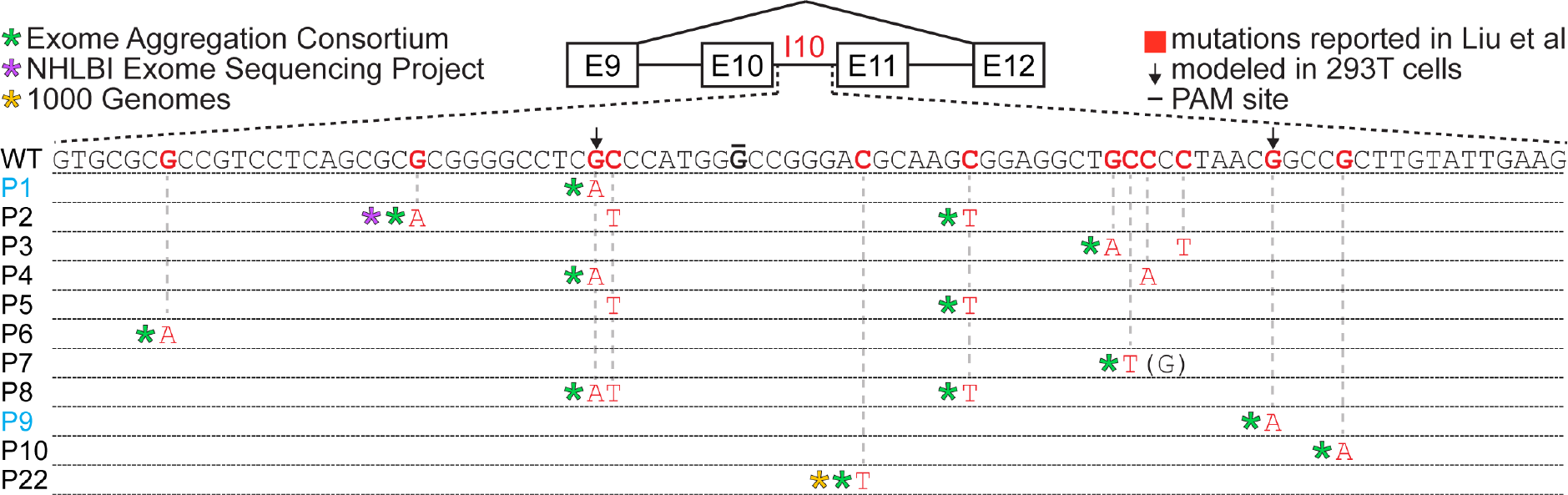
Many reported *UPF1* mutations are identical to genetic variants. Illustration of the mutations in *UPF1* intron 10 (I10) reported by Liu et al. Each row indicates the wildtype (WT) sequence from the reference human genome or mutations reported by Liu et al (P1, patient 1). Blue indicates the mutations that we modeled with genome engineering. Red nucleotides represent positions subject to point mutations reported in Liu et al. Arrows indicate specific mutations that we modeled in 293T cells. The horizontal black line indicates the nucleotide within the protospacer adjacent motif (PAM) site that we mutated to prevent repeated cutting by Cas9. Parentheses indicate where we found genetic variation at a reported mutation position that differed from the specific mutated nucleotide reported by Liu et al. Identical data for the other *UPF1* regions containing mutations reported by Liu et al can be found in **Fig. S2**.

Our discovery that a large fraction of the reported *UPF1* mutations are present in databases of germline genetic variation was surprising for two reasons. First, when strongly cancer-linked mutations occur as germline variants, they frequently manifest as cancer predisposition syndromes. However, no such relationship is known for *UPF1* genetic variants, despite their reportedly high prevalence as identical somatic mutations in PASC. Second, *UPF1* is essential for embryonic viability and development in mammals^15^, zebrafish^16^, and *Drosophila*^17^. As Liu et al reported that all *UPF1* mutations caused missplicing that is expected to disable UPF1 protein function^3^, then those mutations should be incompatible with life when present as inherited genetic variants. Our finding that two reported mutations had no effect on *UPF1* splicing when introduced into their endogenous genomic contexts offers a way to explain this incongruity, at least for the two reported lesions which we studied.

We therefore sought to verify the somatic nature of the *UPF1* mutations described in Liu et al, which was reportedly determined by sequencing both tumors and patient-matched controls. The GenBank accession codes reported in Liu et al corresponded to short nucleotide sequences containing *UPF1* mutations, without corresponding data for patient-matched controls. We contacted the senior author (Dr. YanJun Lu) to request primary sequencing data from patient-matched tumor and normal samples, but neither primary sequencing data from matched samples nor the samples themselves were available.

To further explore whether *UPF1* is recurrently mutated in PASC, we re-analyzed sequencing data from Fang et al^5^ to manually search for *UPF1* mutations (**Tables S3-4**). We focused on the two loci which contained all mutations reported by Liu et al (*UPF1* exons 10 to 11 and exons 21 to 23). Because the relevant introns are very short, they were well-covered by both the whole-exome and whole-genome sequencing used by Fang et al. Using relaxed mutation-calling criteria to maximize sensitivity (details in Methods), we identified somatic *UPF1* mutations in samples from 6 of 17 PASC patients. However, those mutations exhibited genetic characteristics expected of passenger, not driver, mutations. None of those *UPF1* mutations matched the *UPF1* mutations reported by Liu et al, and only one was present at an allelic frequency equal to the allelic frequency of mutant *KRAS*, which is a known driver and which we detected in samples from all PASC patients (median allelic frequencies of 12% versus 34% for *UPF1* versus *KRAS* mutations). Furthermore, we also identified *UPF1* mutations in samples from patients with non-adenosquamous tumors (3 of 34 pancreatic ductal adenocarcinomas), whereas Liu et al reported finding no *UPF1* mutations in non-adenosquamous pancreatic cancers (0 of 29). In concert with Witkiewicz et al^4^ and Hayashi et al’s^6^ reports of finding no *UPF1* mutations in their PASC samples, these analyses suggest that *UPF1* is not a frequent or adenosquamous-specific mutational target in most PASC cohorts.

*UPF1*’s role in the pathogenesis of PASC has been unclear and controversial given the seeming discrepancies between its mutational spectrum in different PASC cohorts. Although it is difficult to conclusively prove that a specific genetic change does not promote cancer, we were unable to detect biological or molecular changes arising from two mutations reported by Liu et al. *UPF1*’s status as an essential gene and our discovery that many reported *UPF1* mutations occur as germline genetic variants of no known pathogenicity together suggest that other *UPF1* mutations reported by Liu et al could similarly represent genetic differences that do not functionally contribute to PASC. Our study highlights the need for continued study of the PASC mutational spectrum in order to understand the molecular basis of this disease.

## ACKNOWLEDGEMENTS

We thank members of the Bradley, Ventura, and Leach laboratories for comments and suggestions. We specifically thank the following individuals for their technical help and support: Olivera Grbovic-Huezo for pancreatic injections, Paul Ogrodowski and Jonathan Bermeo for assistance with mouse work and tissue harvest, Maria S. Jiao and the MSK Center For Comparative Medicine and Pathology Facility for p40 IHC, and Miles Wilkinson for discussing our findings. J.T.P. was supported in part by the NIH/NCI (T32 CA009657). D.U. was supported in part by the NIH/NIGMS (T32 GM007270). R.K. was supported in part by the NIH/NCI (T32 CA160001). S.D.L. was supported in part by the NIH/NCI (R01 CA204228). A.V. was supported in part by the Cycle for Survival’s Equinox Innovation Award in Rare Cancers and a Functional Genomics Initiative grant (AV). R.K.B. is a Scholar of The Leukemia and Lymphoma Society (1344-18).

## AUTHOR CONTRIBUTIONS

J.T.P., R.K., and R.K.B. designed the study. J.T.P. and D.U. performed molecular experiments in human cells. R.K. performed mouse experiments. L.E.-H. assisted with pancreatic injections and performed ultrasound imaging. G.A. analyzed tumor histology and IHC data. A.V. and S.L. contributed to the design of mouse experiments. J.T.P., R.K., and R.K.B. wrote the paper, with input from all authors.

## DATA AVAILABILITY

Whole-genome and whole-exome sequencing data from Fang et al was downloaded from the Sequence Read Archive (accession SRP107982). Genetic variation data was downloaded from the Exome Aggregation Consortium (http://exac.broadinstitute.org/), 1000 Genomes Project (http://www.internationalgenome.org/), and NHLBI Exome Sequencing Project (http://evs.gs.washington.edu/EVS/). Other data that support this study’s findings are available from the authors upon reasonable request.

## SUPPLEMENTAL FILES

**Supplementary Figures**

Supplementary Figures 1-3. Fig. S3 contains uncropped gels related to the main and Supplementary Figures.

**Supplementary Table 1.** Histopathological classification of pancreatic tumors derived from orthotopic injection of control or *Upf1*-targeted KPC cells. Tumors displaying >25% squamous differentiation were classified as positive for squamous features.

**Supplementary Table 2.** Overlap between *UPF1* mutations reported by Liu et al and genetic variants present in the 1000 Genomes, NHLBI Exome Sequencing Project, and ExAC databases. Genomic coordinates are specified with respect to the GRCh37/hg19 genome assembly.

**Supplementary Table 3.** Summary statistics of somatic mutations in *UPF1* identified in our re-analysis of whole-exome and whole-genome sequencing data from Fang et al.

**Supplementary Table 4.** All genetic differences in *UPF1* and *KRAS* from the reference genome, including both somatic mutations and inherited genetic variants, that we identified in our re-analysis of all samples from Fang et al’s cohorts.

**Supplementary Table 5.** Sequences of PCR primers and oligos.

## METHODS

### Construction of mouse KPC cells carrying a deletion of *Upf1* exons 10 and 11

Mouse KPC cells (Kras^G12D^; p53^R172H/null^; Pdx1-Cre) were obtained from Dr. Steven Leach’s lab (Dartmouth College) and were cultured in DMEM (GIBCO) supplemented with 10% fetal bovine serum (FBS) and 1% Penicillin/Streptomycin (GIBCO). All cell lines were incubated at 37°C and 5% CO2. Guide RNAs targeting mouse *Upf1* introns 9 and 11 were cloned into a paired guide expression vector (px333) as previously described^18^. An EcoRI-XhoI fragment containing the double U6-sgRNA cassette and Flag-tagged Cas9 was then ligated into the EcoRI-XhoI-digested pacAd5 shuttle vector. Recombinant adenoviruses were generated by Viraquest (Ad-Upf1 and Ad-Cas9) or purchased from the University of Iowa (Ad-Cre). KPC cells were infected with (5×10^6^ PFU) of Ad-Cas9 or Ad-Upf1 in each well of a 6-well plate.

Genomic DNA was extracted 48 hours post infection to confirm excision of *Upf1* exons 10 and 11. For PCR analysis of genomic DNA, cells were collected in lysis buffer (100 nM Tris-HCl at pH 8.5, 5 mM EDTA, 0.2% SDS, 200 mM NaCl supplemented with fresh proteinase K at a final concentration of 100 ng/mL). Genomic DNA was extracted with phenol-chloroform-isoamylic alcohol and precipitated in ethanol, and the DNA pellet was dried and resuspended in double-distilled water. For RT-PCR, total RNA was extracted with TRIzol (Life Technologies) following the manufacturer’s instructions. cDNA was synthesized using SuperScript III (ThermoFisher) following the manufacturer’s instructions.

### Immunoblots in mouse KPC cells

Cells were lysed in 1X RIPA buffer with protease and phosphatase inhibitors. Fifteen micrograms of protein was separated on 4-10% acrylamide/bisacrylamide gels, transferred onto PVDF membranes, and blocked for 1 hour in 5% milk in 1X TBST. The membranes were incubated with rabbit UPF1 antibody (CST #9435) antibody used at 1:1000 dilution in 5% BSA 1X TBST overnight at 4°C or mouse Tubulin (Sigma T9206) used at 1:2000 dilution in 5% milk 1X TBST for 1 hour at room temperature. Following primary antibody incubation, the membranes were washed three times with 1X TBST buffer at room temperature and probed with rabbit or mouse horseradish peroxidase-linked secondary antibody (1:5000; ECL NA931 mouse, NA934V Rabbit). The Western blot signal was detected using the ECL Prime (RPN322) kit and the blot was exposed to an X-Ray film, which was developed using the Konica Minolta SRX 101A film processor.

### Tumorigenicity and metastasis assays

KPC cells carrying *Upf1* ΔE10-11 or an empty Cas9 control were mixed in 1:1 Matrigel (BD Biosciences) and simple media to a final concentration of 100,000 cells in 30 μL of total volume. Cells were orthotopically implanted into the tails of the pancreata of B6 albino mice (Charles River). Ten mice were implanted with each stable, genetically engineered cell line. Tumor growth was measured weekly via 3D-ultrasound starting at 10 days post-implantation. For survival assessment, animals were sacrificed following the endpoints approved by IACUC: (i) animals showing signs of significant discomfort, (ii) ascites or overt signs of tumor metastasis or gastrointestinal bleeding (blood in stool), (iii) animals losing >15% of their body weight, and (iv) animals with tumors >2 cm in diameter. Investigators responsible for monitoring and measuring the xenografts of individual tumors were not blinded. All animal studies were performed in accordance with institutional and national animal regulations. Animal protocols were approved by the Institutional Animal Care and Use Committee at Memorial Sloan-Kettering Cancer Center (IACUC-MSKCC). Survival curves and statistics were performed using PRISM.

### Immunohistochemistry (IHC) and histopathological analysis

Paraffin sections were dewaxed in xylene and hydrated in graded alcohols. Endogenous peroxidase activity was blocked by immersing the slides in 1% hydrogen peroxide in PBS for 15 minutes. Pretreatment was performed in a steamer using 10 mM citrate buffer (pH 6.0) for 30 minutes. Sections were incubated overnight with a primary rabbit polyclonal antibody against p40-DeltaNp63 (Abcam, ab166857) diluted at a ratio of 1:100. Sections were washed with PBS and incubated with an appropriate secondary antibody followed by avidin-biotin complexes (Vector Laboratories, Burlingame, CA, PK-6100). The antibody reaction was visualized with 3-3’ diaminobenzidine (Sigma, D8001) followed by counterstaining with hematoxylin. Tissue sections were dehydrated in graded alcohols, cleared in xylene, and mounted. For p40 (ΔNp63) IHC, expression was defined based on nuclear labeling.

### Culture and genome engineering of HEK 293T cells

HEK 293T cells were cultured in DMEM media (GIBCO) supplemented with 10% FBS (GIBCO), 100 IU Penicillin, and 100 mg/mL Streptomycin (PenStrep, GIBCO). Cells were cultured at 37°C and 5% CO_2_. Cells were split at a ratio of 1:10 once they reached 90-100% confluency as needed.

Guide RNAs for all cell lines were designed using the GuideScan 1.0 software package^19^ and sequences were chosen among those predicted to have the highest cutting efficiency and specificity scores. DNA oligos for all guide RNAs were synthesized by IDT and amplified using primers that appended homology arms to facilitate ligation into the pX459/Cas9 expression plasmid^20^ by Gibson assembly^21^. Gibson assembly reactions were transformed into NEB Stable Competent *E. coli* and resulting sgRNA expression plasmids were amplified and purified using standard protocols.

Ultramers for homology directed repair (HDR) were designed using previously described strategies^22^ and synthesized by IDT, Inc. HEK 293T cells were transiently transfected with 1 μg/mL sgRNA/pX459 Cas9 expression plasmid and 20 nM HDR Ultramer using Lipofectamine 2000 that was diluted in Opti-MEM Reduced Serum Medium (ThermoFisher). Cells were incubated at 37°C for 24 hours, at which time the transfection medium was replaced with DMEM media (GIBCO) supplemented with 10% FBS (GIBCO), 100 IU Penicillin, 100 mg/mL Streptomycin (PenStrep, GIBCO), and 2 mg/mL puromycin. Cells were then incubated for 48-72 hours in DMEM media (GIBCO) supplemented with 10% FBS (GIBCO), 100 IU Penicillin, 100 mg/mL Streptomycin (PenStrep, GIBCO), and 2 mg/mL puromycin, at which time genomic DNA was extracted and regions of interest were amplified using the appropriate oligos (**Table S5**).

Genome engineering was validated using genomic DNA as follows. Amplicons from genomic DNA PCR were ligated into vectors using the Zero Blunt TOPO PCR cloning system (ThermoFisher) and the presence of the desired mutations was validated using Sanger sequencing (GENEWIZ). Polyclonal cell populations were then diluted and sorted into 96-well plates using a BD FACS Aria II flow cytometer (BD Biosciences), such that each well contained on average one cell, which were grown in DMEM media (GIBCO) supplemented with 20% FBS (GIBCO), 100 IU Penicillin, and 100 mg/mL Streptomycin (PenStrep, GIBCO). Once cells in 96-well plates reached confluency, they were transferred to 24-well plates and allowed to grow to confluency for genomic DNA extraction and Sanger sequencing.

### NMD efficiency measurement

NMD efficiency was estimated using the beta-globin reporter system^11^. HEK 293T cells engineered with the reported mutations were plated at 10-15% confluency on pol-L-lysine coated 12-well plates. Cells were co-transfected 24 hours later with 1 μg of phCMV-MUP plasmid (transfection control) and 1 μg of either pmCMV-Gl-Norm (normal termination codon) or pmCMV-Gl-39Ter (premature termination codon) using Lipofectamine 3000 (Invitrogen) according to the manufacturer’s protocol. After 48 hours the cells were close to confluency, at which time they were lysed using 1 mL of Trizol Reagent (Invitrogen) per well. The lysate was collected, and total RNA was extracted according to the manufacturer’s protocol. The RNA was further purified and DNase treated using the Direct-zol RNA MiniPrep Kit (Zymo Research) according to the manufacturer’s protocol.

Residual plasmid DNA was removed using DNaseI (Amplification Grade, Invitrogen) according to the manufacturer’s protocol from 600 ng of the extracted RNA. cDNA synthesis was then performed using SuperScript IV Reverse Transcriptase (Invitrogen) with oligo dT primers according to the manufacturer’s protocol. The cDNA synthesis reaction was diluted 1:50 and 4 μL was used for a 10 μL qPCR reaction with PowerUp™ SYBR™ Green Master Mix (ThermoFisher) and primers specific for the reporter mRNA (**Table S5**) diluted to a working concentration of 100 nM for each primer. qPCR reactions were performed in technical triplicate for three different biological replicates in 384-well plates (ThermoFisher) using an ABI QuantStudio 5 Real-Time PCR System (ThermoFisher). The levels of pmCMV-Gl-Norm and pmCMV-Gl-39Ter cDNA were normalized to phCMV-MUP mRNA abundance for each sample, and levels of pmCMV-Gl-39Ter cDNA were plotted relative to levels of pmCMV-Gl-Norm for each replicate in each cell type.

### Immunoblots in HEK 293T cells

Cells were lysed using a buffer containing 150 mM NaCl, 1% NP-40 and 50 mM Tris pH 8.0 supplemented with a phosphatase inhibitor (ThermoFisher) and protease inhibitor (ThermoFisher). After the lysis buffer was added, cells were frozen at −80°C and thawed for three cycles, then incubated on ice for 15 minutes. The lysed cells were centrifuged at 10,000 x g for 15 minutes, and the supernatant was collected to determine total protein concentration. Total protein concentration was determined using a Q-bit (ThermoFisher) and 20 μg of total protein was used for electrophoresis. Following electrophoresis, protein was transferred to nitrocellulose membrane (Novex) in transfer buffer containing 10% methanol overnight at 4°C. The blot was blocked with Odyssey^®^ Blocking Buffer (LI-COR Biosciences) for 1 hour at room temperature and probed using a 1:1000 dilution of 0.514 mg/mL UPF1 antibody (Abcam, 109363) overnight with shaking at 4°C. Following overnight incubation, the blot was washed three times with 1X TBST buffer at room temperature and probed with rabbit secondary antibody (IRDye 680RD goat anti-rabbit) for 1 hour at room temperature. The blot was then imaged and UPF1 abundance was quantified using band intensity in Fiji (v2.0.0).

### Isoform detection by RT-PCR in HEK 293T cell lines

Total RNA was extracted using the RNeasy Plus Mini Kit (Qiagen). cDNA was synthesized using oligo dT primers and SuperScript IV Reverse Transcriptase (ThermoFisher) following the manufacturer’s protocol. PCR was carried out with primers targeting *UPF1* exons 9 and 12 using Q5® High-Fidelity DNA Polymerase (NEB) (**Table S5**). PCR products were run in a 2% agarose slab gel and stained with ethidium bromide for visualization by UV shadowing (Bio-Rad Molecular Imager Gel Doc XR+).

A positive control for the amplification of the truncated *UPF1* variant missing exons 10 and 11 was synthesized as a double stranded DNA gBlock (IDT). Two different amounts (10 fg and 1 fg) of this “spike in” control was added to separate PCRs containing cDNA from wild-type HEK 293T cells and amplified in the same manner as described above.

To quantify the degree of exon skipping (percent ΔE10-11), a background subtraction across the entire gel was first performed in Fiji using a rolling ball radius of 100 pixels. Next, the integrated density of each band was determined, and the density of the lower band in each lane was divided by the total density in that same lane by summing the integrated densities of the upper and lower bands.

### Re-analysis of genomic DNA sequencing data from Fang et al

Whole-genome and whole-exome sequencing data from Fang et al^5^ was downloaded from the Sequence Read Archive (accession number SRP107982) and mapped to the *UPF1* (chr19:18940305-18979266; hg19/GRCh37 assembly) and *KRAS* (chr12:25356390-25405419; hg19/GRCh37 assembly) gene loci. The first 10 nt of all reads were trimmed off due to low sequencing quality and the trimmed reads were mapped with Bowtie v1.0.0^23^ with the arguments ‘-v 3 -k 1 -m 1 --best --strata --minins 0 --maxins 1000 --fr’. Mapped reads were visualized in IGV^24^. Mutation/genetic variant calling thresholds (read coverage depth ≥9 reads and ≥2 reads supporting the mutation/variant) were chosen in order to allow detection of a hotspot *KRAS* mutation (G12 or G13) in every PASC sample in the cohort. That criteria ensured that our thresholds were appropriate for discovering known cancer driver mutations in all samples. A genetic difference from the reference genome was defined as a somatic mutation if it was called in a tumor sample but not in the corresponding patient-matched normal control sample.

