## Supplementary Figures for "The origins and consequences of *UPF1* variants in pancreatic adenosquamous carcinoma"

**SUPPLEMENTARY FIGURE 1**

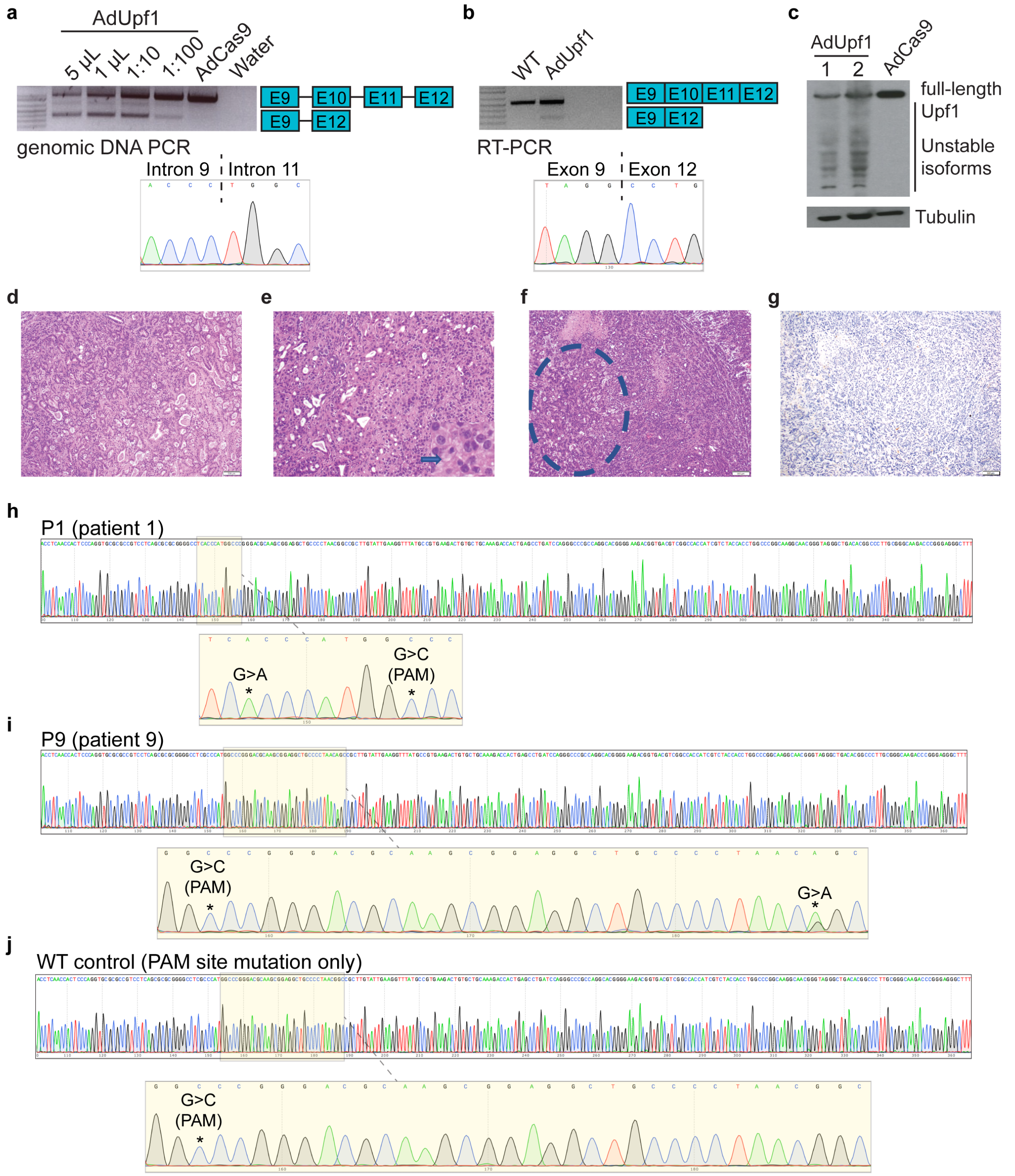


**Supplementary Figure 1. Validation experiments in mouse KPC and human 293T cells.**
(**a**) Titration of adenovirus expressing Cas9 only (AdCas9) or Cas9 and guide RNAs targeting intron 9 and intron 11 of mouse *Upf1* (AdUpf1) in KPC cells. Gel corresponds to PCR using genomic DNA with primers in exons 9 and 12 (E9, E12). The lower band, corresponding to excision of exons 10 and 11, was

only observed in AdUpf1 conditions. This band was excised and sequence-verified by Sanger sequencing (sequence trace displayed below).
(**b**) As (a), but assaying a cDNA library instead of genomic DNA. The lower band, corresponding to spliced mRNA lacking exons 10 and 11, was only observed in AdUpf1 conditions.
(**c**) Immunoblot with a probe against UPF1. Levels of full-length UPF1 protein are lower in AdUpf1 conditions, as expected. 1 and 2 indicate biological replicates. Tubulin serves as a loading control.
(**d**) Representative hematoxylin and eosin (H&E) staining of a pancreatic tumor resulting from orthotopic injection of control KPC cells displaying features of a moderately to poorly differentiated pancreatic ductal adenocarcinoma. Tumors were composed of medium-size duct-like structures and small tubular glands with lower mucin production.
(**e**) Representative H&E image illustrating a moderately to poorly differentiated pancreatic ductal adenocarcinoma resulting from orthotopic injection of *Upf1*-targeted KPC cells. Depicted here is a section of the poorly differentiated component (arrow), which was characterized by solid sheets of tumor cells with large eosinophilic cytoplasms and marked nuclear polymorphism.
(**f**) Representative H&E image of a pancreatic tumor resulting from orthotopic injection of *Upf1*-targeted KPC cells. The dashed circle marks a moderately differentiated component; the remainder is poorly differentiated.
(**g**) Representative IHC image of a pancreatic tumor resulting from orthotopic injection of *Upf1*-targeted KPC cells for the squamous marker p40 (ΔNp63). No expression of the marker was observed in tumor cells.
(**h**) Sanger sequencing of genomic DNA from engineered 293T cells verifying introduction of the mutation IVS10+31G>A in a homozygous state, as reported by Liu et al for patient P1, as well as a single PAM site mutation.
(**i**) Sanger sequencing of genomic DNA from engineered 293T cells verifying introduction of the IVS10-17G>A mutation in a heterozygous state, as reported by Liu et al for patient P9, as well as a single PAM site mutation.
(**j**) Sanger sequencing of genomic DNA from engineered 293T cells verifying introduction of a single PAM site mutation as a wild-type control.
See **Fig. S3** for uncropped gels.

### SUPPLEMENTARY FIGURE 2

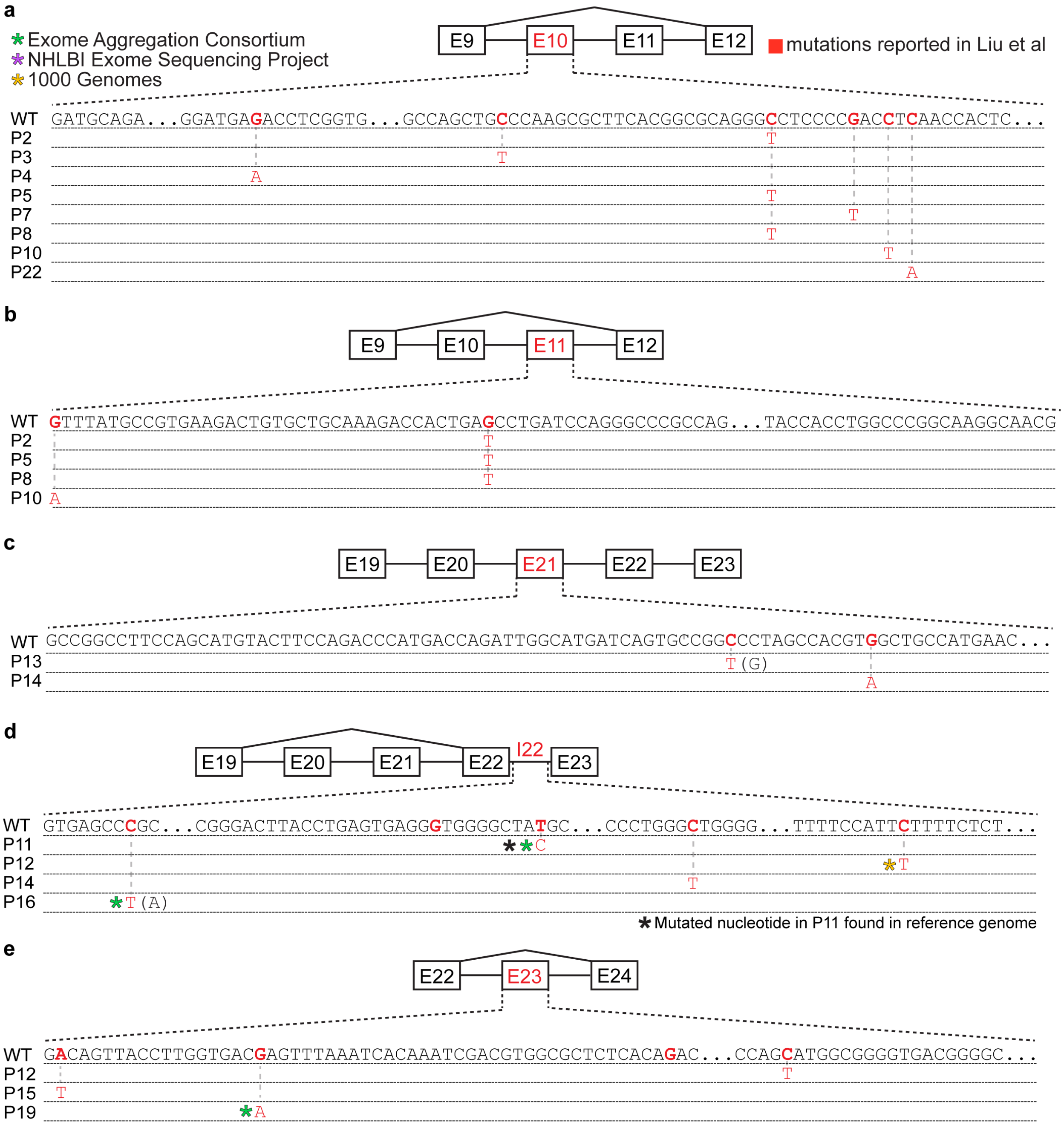


**Supplementary Figure 2. Many reported *UPF1* mutations are identical to genetic variants.** Illustration of the mutations in *UPF1* (**a**) exon 10, (**b**) exon 11, (**c**) exon 21, (**d**) intron 22, and (**e**) exon 23 reported by Liu et al. Each row indicates the wild-type (WT) sequence from the reference human genome or mutations reported by Liu et al (P1, patient 1). Red nucleotides represent positions subject to point mutations reported in Liu et al. Parentheses indicate where we found genetic variation at a reported mutation position that differed from the specific mutated nucleotide reported by Liu et al. Identical data for intron 10 can be found in **Fig. 2**.

### SUPPLEMENTARY FIGURE 3

Figure 1f
Western blot to detect UPF1 in HEK 293T cells

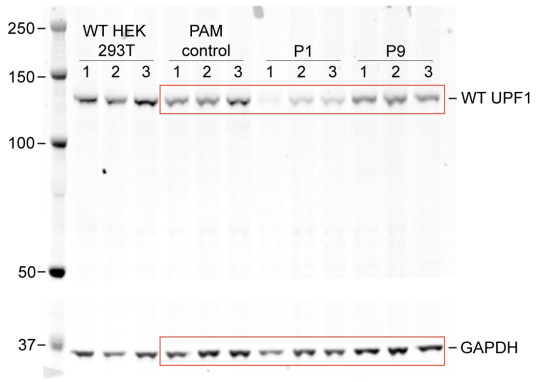


Membrane was cut between the 50 kD and 37 kD bands
The ladder was Precision Plus Protein™ Kaleidoscope™ (BIO-RAD)

Figure 1g
RT-PCR to quantify exon skipping in HEK 293T cells

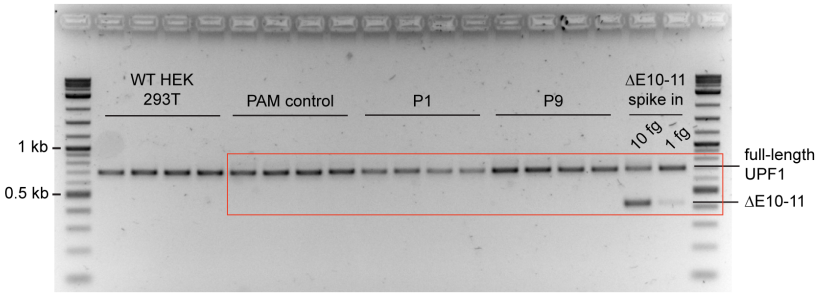


Supplementary Figure 1c
Western blot for detecting UPF1

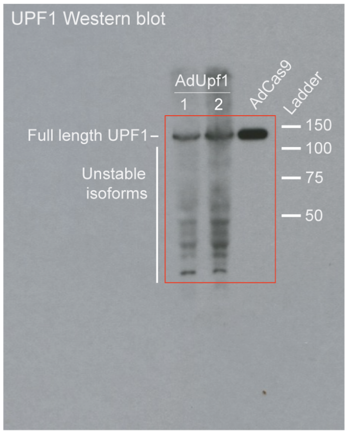


Supplementary Figure 1a
Genomic DNA PCR for *Upf1* exons 9-12

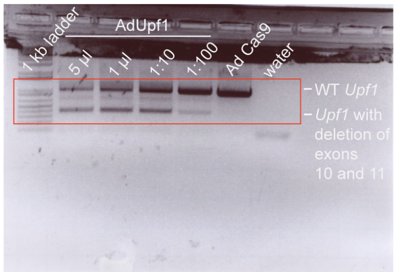


Supplementary Figure 1b
RT-PCR to detect *Upf1* isoform with deletion of exons 10 and 11

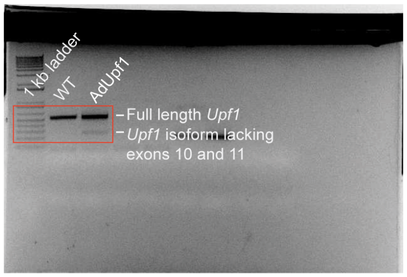


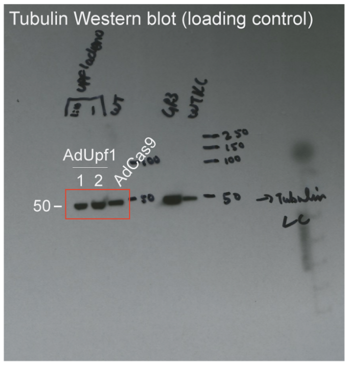


**Supplementary Figure 3. Images of uncropped gels.**
